# The ALS-associated TDP-43^M337V^ mutation dysregulates microglia-derived extracellular microRNAs in a sex-specific manner

**DOI:** 10.1101/2023.11.30.569424

**Authors:** Eleni Christoforidou, Libby Moody, Greig Joilin, Fabio A. Simoes, David Gordon, Kevin Talbot, Majid Hafezparast

## Abstract

Evidence suggests the presence of microglial activation and microRNA (miRNA) dysregulation in amyotrophic lateral sclerosis (ALS), the most common form of adult motor neuron disease. However, few studies have investigated whether the miRNA dysregulation may originate from microglia. Furthermore, TDP-43, involved in miRNA biogenesis, aggregates in tissues of ∼98% of ALS cases. Thus, this study aimed to determine whether expression of the ALS-linked TDP-43^M337V^ mutation in a transgenic mouse model dysregulates microglia-derived miRNAs. RNA sequencing identified several dysregulated miRNAs released by transgenic microglia, and a differential miRNA release by lipopolysaccharide-stimulated microglia, which was more pronounced in cells from female mice. We validated the downregulation of two candidate miRNAs, miR-16-5p and miR-99a-5p by reverse transcriptase-quantitative polymerase chain reaction (RT-qPCR), and identified their predicted targets, which include primarily genes involved in neuronal development and function. These results suggest that altered TDP-43 function leads to changes in the miRNA population released by microglia in a sex dependent manner, which may in turn influence disease progression in ALS. This has important implications for the role of neuroinflammation in ALS pathology and could provide potential therapeutic targets.

## Introduction

Amyotrophic lateral sclerosis (ALS) is the most severe and most common form of motor neuron degeneration in adults, caused by the death of motor neurons in the motor cortex, brainstem, and spinal cord. Although ALS was traditionally considered as having cell-autonomous mechanisms (i.e., damage within the motor neurons being enough to cause disease), numerous lines of evidence suggest that the death of motor neurons is influenced by non-neuronal cells such as astrocytes and microglia (Boillée et al., 2006, Kang et al., 2013, Yamanaka and Komine, 2018, Vahsen et al., 2021), and non-cell-autonomous mechanisms appear to play significant roles in the disease onset and/or progression. In fact, studies on superoxide dismutase 1 (SOD1) transgenic mouse models suggest that isolated expression of disease-associated protein variants in motor neurons appeared to be insufficient for disease onset (Lino et al., 2002, Pramatarova et al., 2001), which requires simultaneous expression within glial cells (Clement et al., 2003, Yamanaka et al., 2008). In line with this, there is further evidence that the expression of SOD1^G37R^ in motor neurons underlies disease onset and that reducing the expression of the mutant protein within microglia slows disease progression (Boillée et al., 2006). Importantly, increased numbers of activated microglia have been observed in the central nervous system (CNS) of ALS mouse models and patients (Hall et al., 1998, McGeer et al., 1993). Specifically, studies on post-mortem tissues have shown increased levels of activated microglia in areas of the brain with neuronal loss (Boillée et al., 2006, Polazzi and Monti, 2010). Most recently, microglia derived from induced pluripotent stem cells from ALS patients carrying the C9orf72 mutation have been shown to exert toxicity on motor neurons via pro-inflammatory pathways (Vahsen et al., 2023).

MicroRNAs (miRNAs) are small non-coding RNAs (ncRNAs) which bind to complementary messenger RNA (mRNA) sequences, resulting in gene silencing via degradation or translational repression (Bartel, 2009). Evidence for a role for miRNAs in ALS pathology comes from the observation of a differential miRNA expression profile between ALS patients and healthy controls in circulating fluids, particularly in the blood (Joilin et al., 2019, Joilin et al., 2020, Joilin et al., 2022), giving rise to the opportunity of using them as potential biomarkers (Joilin et al., 2019, Ricci et al., 2018). However, the source of these miRNAs is unknown, and it is unclear whether these are released by atrophied muscles, degenerating motor neurons, or other cell types such as the glia. In addition, FUS and TDP-43, proteins associated with ALS, are directly involved in miRNA processing (Morlando et al., 2012, Kawahara and Mieda-Sato, 2012). Consequently, their mislocalisation in cytoplasmic aggregates in ALS may be associated with the miRNA dysregulation observed in ALS patients.

A recently described transgenic mouse model carrying the human *TARDBP* gene (encoding TDP-43) with the ALS-associated *M337V* mutation has provided new insights into the mechanisms of TDP-43 pathology (Gordon et al., 2019). This mouse model expresses a single copy of the gene integrated into a defined neutral position within the mouse genome. Expression of this mutant transgene results in a progressive motor deficit, a loss of neuromuscular junction integrity, and a decrease in survival. Furthermore, primary motor neurons from this model exhibit a defect in stress granule dynamics and TDP-43 mislocalisation. Given the known role of TDP-43 in RNA metabolism and miRNA processing, this model is useful in facilitating the investigation of altered miRNA processes in ALS.

We previously reviewed the evidence implicating reactive microglia and dysregulated miRNAs in ALS, and explored how microglia may potentially be the source of this miRNA dysregulation (Christoforidou et al., 2020). We concluded that this is likely because microglia release extracellular vesicles containing miRNAs, which play a role in gene regulation, and alterations in these miRNAs, along with inflammation and changes in microglial phenotypes, are observed in ALS patients. Based on those observations, here we hypothesised that expression of the ALS-linked TDP-43^M337V^ mutation dysregulates microglia-derived miRNAs and that this may further be affected by their activation state. To test this hypothesis, we used next-generation sequencing to profile and interrogate the comparative expression of miRNAs released by transgenic and non-transgenic microglia, in the presence or absence of the pro-inflammatory stimulus, lipopolysaccharide (LPS).

Here, we found that the TARDBP^M337V^ transgene did not significantly affect the microglial response to LPS in terms of TNF and IL-1β cytokine induction. However, the transgene appears to influence miRNA release in a sex-dependent manner, with female microglia showing a more pronounced effect. We identified a total of 391 miRNAs, with males and females showing distinct profiles of upregulated and downregulated miRNAs in response to LPS treatment. Furthermore, genotype significantly impacted miRNA release only in female samples, implying the presence of the TDP-43 transgene influences miRNA release, particularly in females. Notably, this effect seemed to be dose-dependent, with greater miRNA dysregulation in homozygous lines. These findings reveal a complex interplay between genotype, treatment response, and sex in determining the miRNA profile released from microglia, potentially influencing distinct mechanisms of neuroinflammation.

## Results

### Impact of the TARDBP^M337V^ transgene on the microglial cytokine response to LPS

Primary mouse microglia from neonatal transgenic (*TARDBP^−/+^*, *TARDBP^−/M337V^*, *TARDBP^M337V/M337V^*) and non-transgenic (*TARDBP^−/−^*) mice of both sexes were stimulated with 250 ng ml^-1^ LPS or vehicle for 24 hours, before the culture medium and cells were collected. To confirm the successful induction of a response in the cells by the LPS stimulus, the gene expression of the pro-inflammatory cytokines TNF and IL-1β was quantified by RT-qPCR. As expected (Diz-Chaves et al., 2012, Kang et al., 2019), there was a significant increase in the expression of both *Tnf* and *Il1b* in all the samples following LPS treatment, compared to vehicle treatment (Fig. 1A-D), regardless of genotype, indicating a successful pro-inflammatory response after 24 hours (main effect of treatment, P<0.0001 for both sexes). Additionally, in female samples, the baseline expression of *Tnf* and *Il1b* in the absence of LPS stimulation (i.e., only vehicle treatment) was not significantly different between cells of different genotype, suggesting that the presence of the human transgene does not alter the baseline expression levels of these pro-inflammatory cytokines (Fig. 1A-D; main effect of genotype, P≥0.05). However, in males, there was a significant main effect of genotype for *Tnf* only (P=0.0090), and the post-hoc analysis revealed this to be due to a significant downregulation of *Tnf* in vehicle-treated *TARDBP^M337V/M337V^*samples, when compared to *TARDBP^−/M337V^* samples (P=0.0143; Fig. 1A), thus indicating a potential influence of transgene copy number; but also due to a significant downregulation of *Tnf* in *TARDBP^M337V/M337V^* samples, when compared to *TARDBP^−/−^* samples (P=0.0251; Fig. 1A), thus indicating an effect of the mutation itself. Finally, for *Tnf*, there was no significant interaction between treatment and genotype in either sex. However, for *Il1b*, there was a statistically significant (P=0.0254) interaction between treatment and genotype in females only. Samples from individual mice (female only) also exhibited significant variation (P<0.0001) in their expression of *Il1b*.

**Figure 1.**
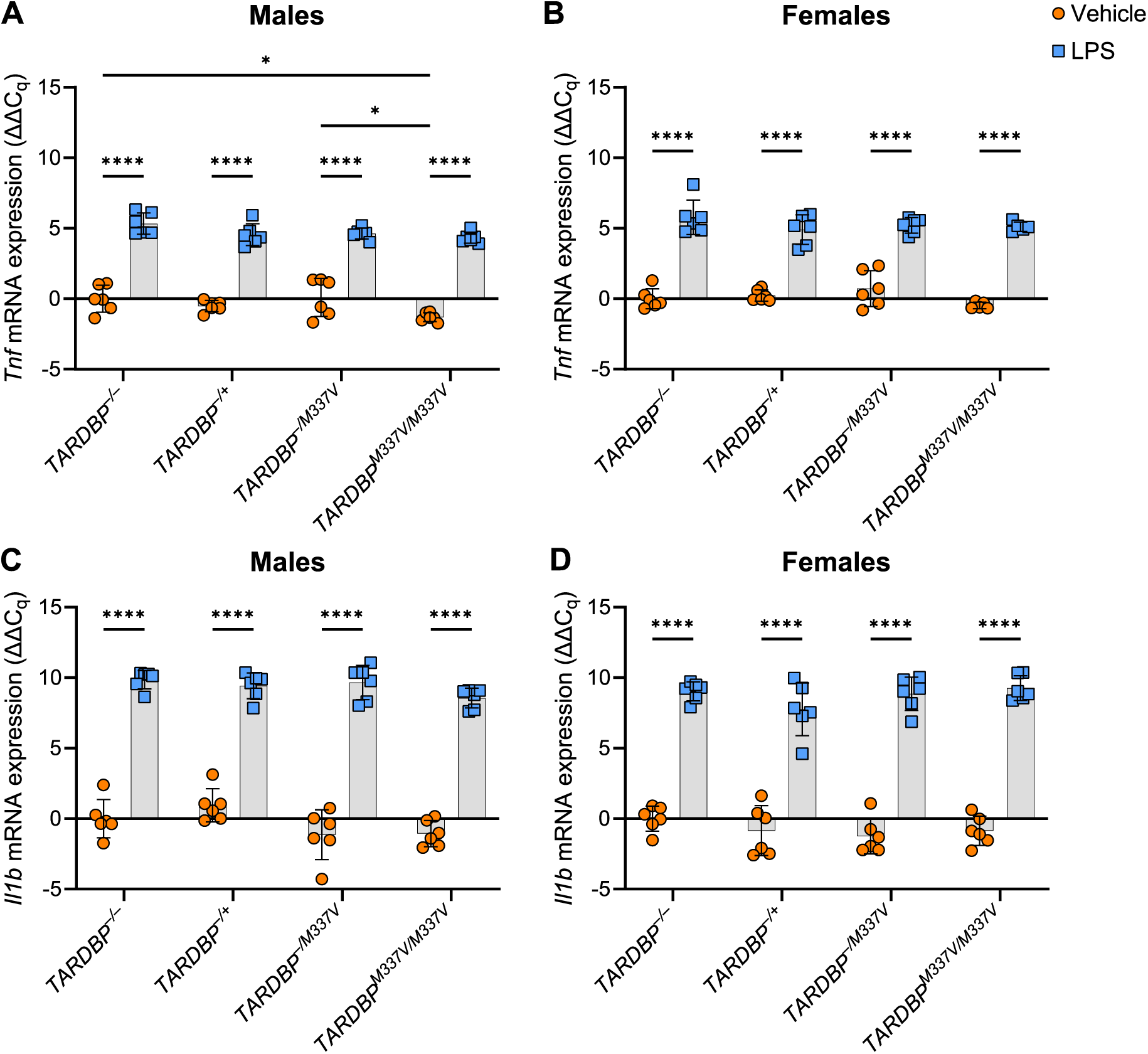
Induction of *Tnf* and *Il1b* expression upon LPS treatment. (**A-B**) *Tnf* mRNA expression. One outlier was removed from the female *TARDBP^M337V/M337V^* data. (**C-D**) *Il1b* mRNA expression. Expression values are normalised to the mean of the vehicle-treated samples from *TARDBP^−/−^* animals within each sex group, following normalisation to housekeeping genes (*Gapdh* and *Pgk1*). Data is shown as mean ± SD. Two-way ANOVA with Šidák’s post-hoc. Main effect of treatment (panels **A**-**D**): P<0.0001. Main effect of genotype (panel **A**): P=0.0090. Genotype × Treatment interaction (panel **D**): P = 0.0254. Pairwise comparisons: *P<0.05, ****P<0.0001. N = 6 biological replicates per genotype per sex.

In summary, these data demonstrate that while the *TARDBP^M337V^*transgene may influence the baseline expression levels of *Tnf* in male microglia, it does not appear to affect the overall response of microglia to LPS stimulation. However, the significant interaction between treatment and genotype in female microglia for *Il1b* indicates that the relationship between transgene presence and cytokine response may be more complex and warrants further investigation. These findings provide an important foundation for understanding how TDP-43 mutations may modulate microglial activation and inflammatory response in the context of ALS.

### Quality of sequencing and alignment information

Following exposure of microglia to LPS or vehicle for 24 hours, the culture medium was collected, and the total RNA released by the cells was extracted. This includes small ncRNAs such as miRNAs released by the microglia during the 24-hour treatment period. This extracellular RNA was used to prepare miRNA libraries for next-generation sequencing. Quality control of the sequencing data revealed that all the per-base sequence qualities (Fig. 2A) and per-sequence quality scores (Fig. 2B) were above the threshold of 28, indicating that the sequencing results of all samples were of good quality. Moreover, the average GC content per read was 54% (Fig. 2C), and as expected, most reads fell at ∼22 bp (corresponding to the average size of miRNAs) and within the ∼24-30 bp range [corresponding to the size range of PIWI-interacting RNA (piRNA); Fig. 2D]. Of the total sequencing reads in each sample, 55% on average mapped to the mouse genome (Fig. 2E), and most of the remaining reads did not map due to being too short (data not shown). Mapped reads mainly consisted of miRNA (33.58%) and ‘other RNA’ (45.32%), which included long non-coding RNA, small nuclear RNA, small nucleolar RNA, and predicted RNA sequences (Fig. 2F). Other constituents of the mapped reads were transfer RNA (tRNA, 8.25%), ribosomal RNA (rRNA, 4.40%), piRNA (0.72%), mRNA (0.70%), hairpin RNA (0.05%) and ‘not characterised’ reads (6.98%) that aligned to locations of the genome that do not correspond to currently known RNA sequences (Fig. 2F).

**Figure 2.**
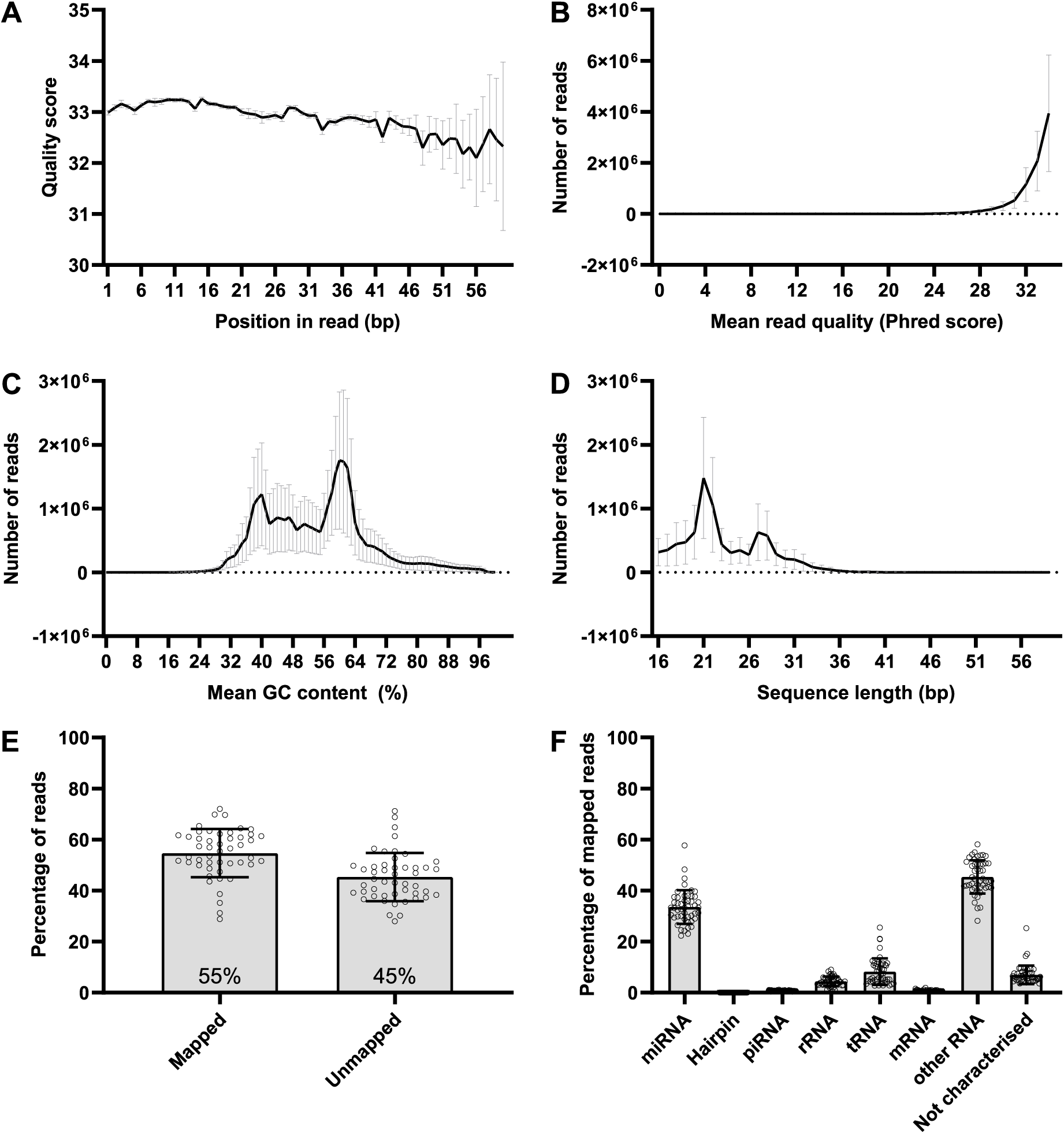
Quality control of sequencing reads and mapping information. (**A**) Per-base sequence quality. (**B**) Per-sequence quality scores. (**C**) Per sequence GC content. (**D**) Sequence length distribution. (**E**) Percentage of mapped and unmapped reads in each sample sequenced (mean percentage value is indicated within the bars). (**F**) Percentage of mapped reads. ‘Not characterised reads are those that aligned to the genome but in a location that does not correspond to currently known RNA sequences. Data is shown as mean ± SD. N = 48 samples.

### The TARDBP^M337V^ transgene dysregulates miRNAs released from microglia in a sex-specific manner

A total of 391 miRNAs were detected in at least one of the samples sequenced. The differential expression analysis of these miRNAs was undertaken separately for males and females. In male samples, after controlling for the genotype effect, we identified three upregulated and one downregulated miRNAs upon LPS treatment, compared to vehicle treatment. On the other hand, in female samples we identified 22 upregulated and 13 downregulated miRNAs (Fig. 3; Table S1), only two of which were common with the males. This stark disparity underscores a potent sex-specific effect on the microglial response to LPS.

**Figure 3.**
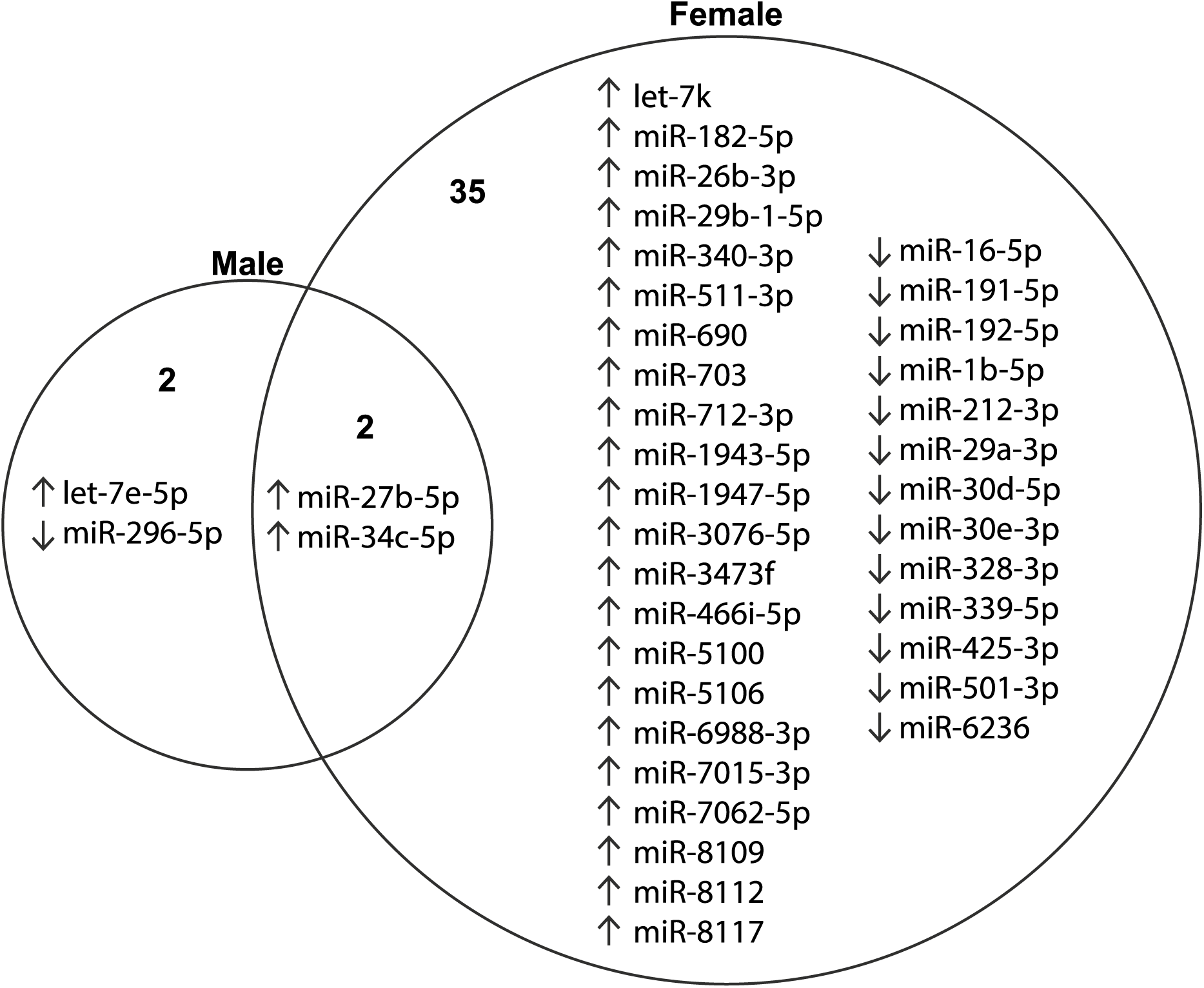
Dysregulated release of miRNAs upon LPS stimulation, after adjusting for genotype effects. ‘Up’ arrows indicate upregulation, whereas ‘Down’ arrows indicate downregulation, when compared to vehicle treatment. N = 3 biological replicates per genotype per sex. A list of these dysregulated miRNAs and associated significance values is also provided in Table S1.

To assess the impact of human transgene expression on extracellular miRNA release from microglia, after adjusting for treatment effects, we compared each transgenic group with the non-transgenic controls, stratified by sex. Notably, genotype significantly affected miRNA release only in female samples (Fig. 4; Table S2). Specifically, we identified two upregulated miRNAs in *TARDBP^−/+^*samples, five in *TARDBP^−/M337V^* samples, and 12 in *TARDBP^M337V/M337V^*samples, compared to non-transgenic controls. Interestingly, the two upregulated miRNAs (miR-9-3p and miR-877-5p) in *TARDBP^−/+^* samples were also found to be upregulated in *TARDBP^−/M337V^*and *TARDBP^M337V/M337V^* samples. The recurrence of these two dysregulated miRNAs across all transgenic genotypes suggests that the presence of the human TDP-43 transgene, irrespective of its mutational status, has a baseline influence. However, we observed a heightened severity in the dysregulation of these two miRNAs in the two mutant transgenic groups compared to the wildtype transgenic group (Table 1A). This pattern suggests that the TDP-43 mutation not only determines the subset of affected miRNAs but also intensifies the degree of their dysregulation, underscoring the mutation’s exacerbating role in perturbing miRNA expression profiles. Furthermore, these two miRNAs showed higher dysregulation in homozygous than hemizygous mutant transgenics (Table 1A), and similarly, there were an additional three miRNAs commonly dysregulated between *TARDBP^−/M337V^* and *TARDBP^M337V/M337V^* samples, but with a higher dysregulation in homozygous than in hemizygous mutant transgenics (Fig. 4; Table 1B). This suggests a dose-dependent effect of the TDP-43 mutation on miRNA release, with higher mutation levels associated with a greater degree of miRNA dysregulation. Overall, the minimal miRNA dysregulation observed in the female wildtype transgenic samples suggests that the presence of the human TDP-43 transgene *per se* exerts a limited impact on miRNA release from microglia, whereas the mutant TDP-43 transgene exerts a larger effect.

**Table 1.**
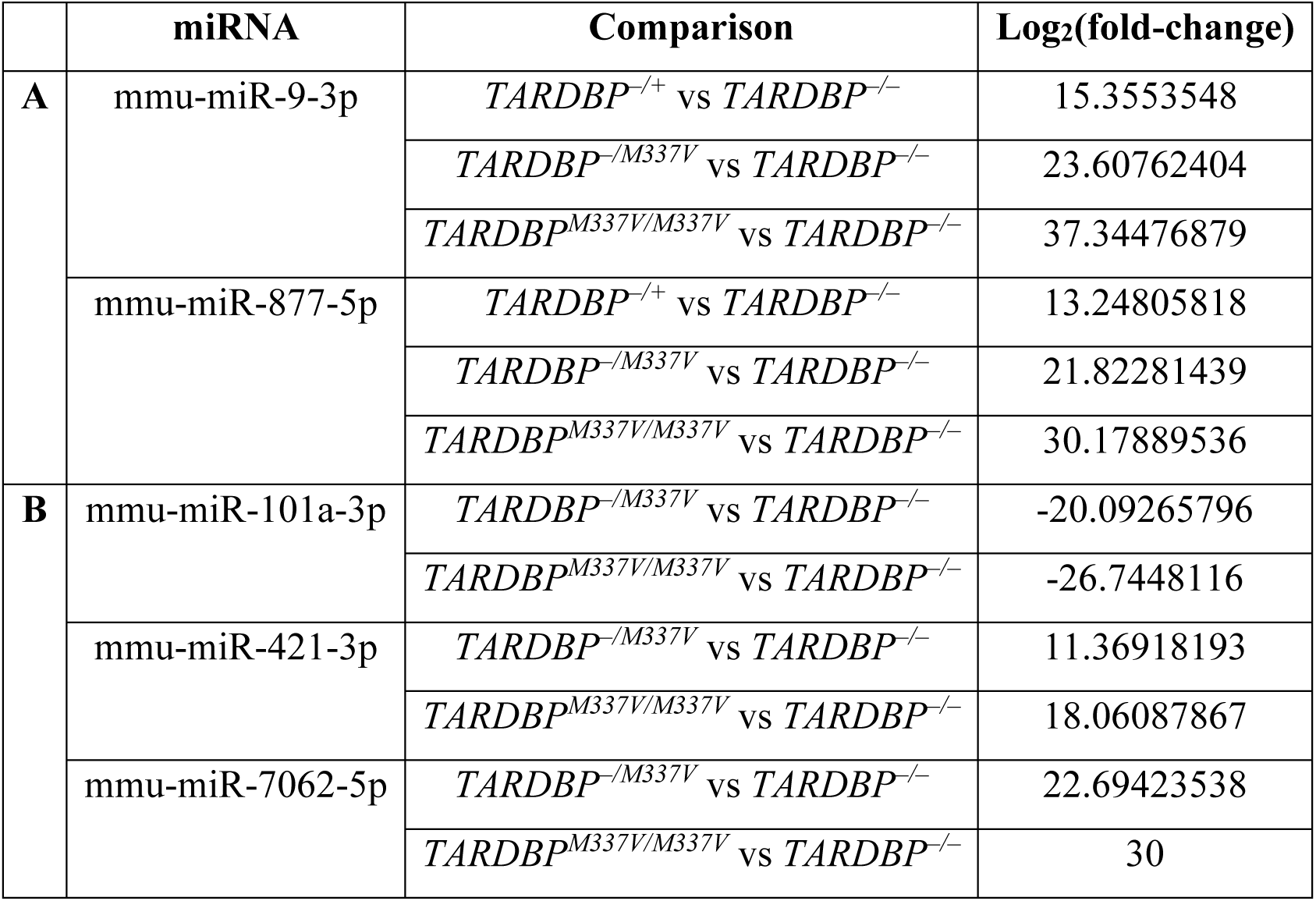
MiRNAs commonly dysregulated in all three female transgenic groups, compared to non-transgenic controls, after adjusting for treatment effects.

**Figure 4.**
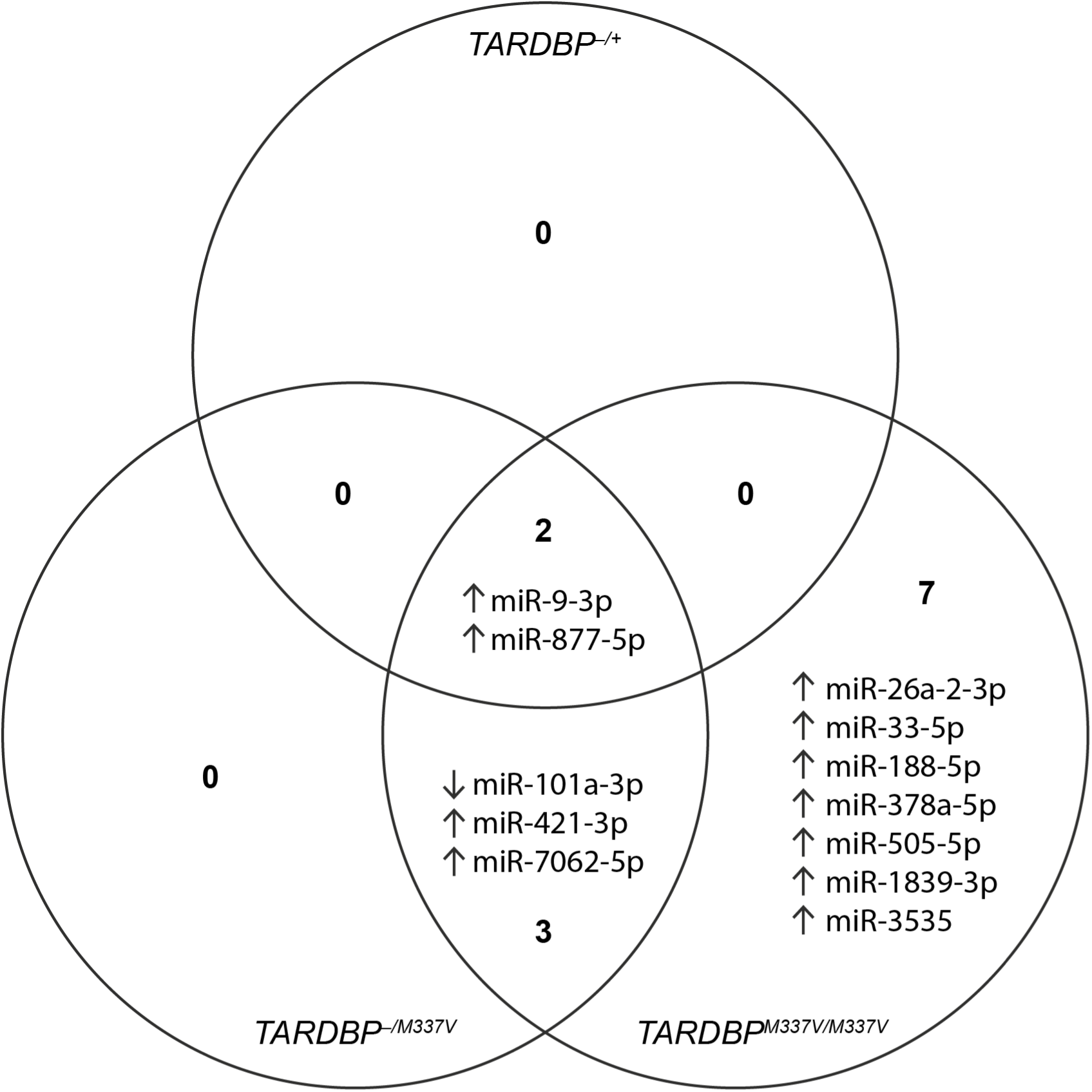
Dysregulated release of miRNAs in female transgenic samples, after adjusting for treatment effects. ‘Up’ arrows indicate upregulation, whereas ‘Down’ arrows indicate downregulation, when compared to non-transgenic controls. N = 3 biological replicates per genotype per sex. A list of these dysregulated miRNAs and associated significance values is also provided in Table S2.

Next, in order to discern how miRNA release following stimulation with LPS differs between genotypes, we examined the interaction between genotype and treatment. We conducted this analysis separately for each sex. Our findings revealed statistically significant miRNA dysregulation for most comparisons and for both sexes (Table S3). There were significant differences in the response to LPS treatment (compared to vehicle treatment), with little overlap between different genotypes (Table S4). Importantly, for each genotype pairwise comparison, different miRNAs were dysregulated in males than in females, with minimal overlap between the sexes (Fig. 5). Like the genotype-only effects discussed earlier, when juxtaposing the homozygous mutant transgenics against the hemizygous wildtype transgenics, there was a relatively modest degree of dysregulation (one dysregulated miRNA in each sex). Conversely, the comparison of the homozygous mutant transgenics to the non-transgenics showcases a greater number of dysregulated miRNAs (11 dysregulated miRNAs in each sex; Fig. 5). Given this pronounced effect, our subsequent RT-qPCR validation was directed towards those miRNAs identified as dysregulated in the comparison between homozygous mutant transgenics and non-transgenics. In this way we aimed to provide a deeper insight into the most consequential alterations in miRNA profiles, underpinned by the presence of the TDP-43 mutation.

**Figure 5.**
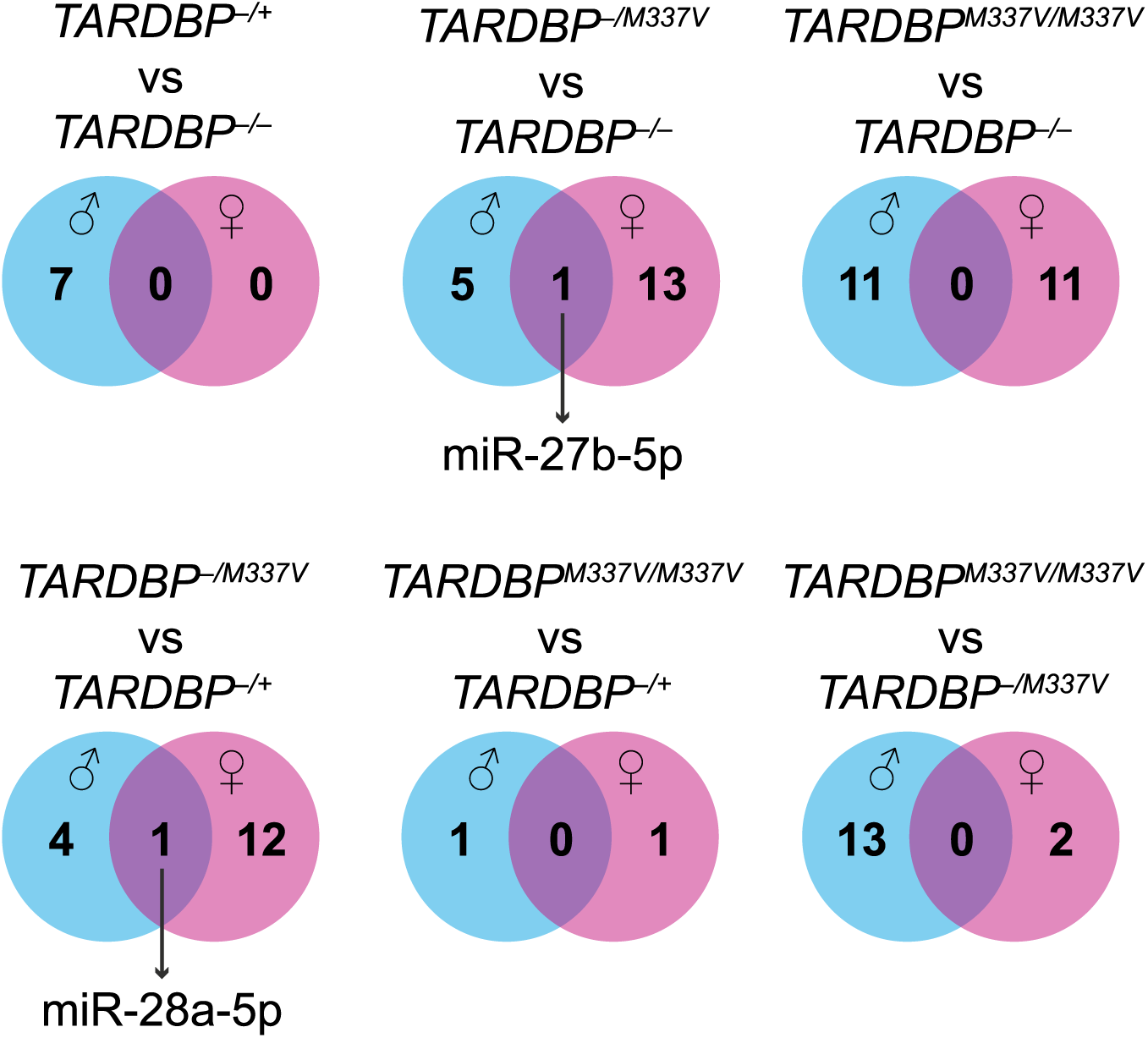
Comparison of dysregulated release of miRNAs upon LPS treatment between male and female samples. N = 3 biological replicates per genotype per sex. A list of the dysregulated miRNAs and associated significance values is provided in Table S3.

Overall, our findings underscore the differential response of male and female microglia to LPS treatment and demonstrate the significant impact of the presence and dose of the TDP-43 mutation on miRNA dysregulation, particularly in female microglia. These disparities in miRNA release profiles could potentially drive distinct molecular mechanisms of neuroinflammation in males and females, as well as in different genotypes.

### Validation of sequencing results by RT-qPCR

Next, we used RT-qPCR to quantify the levels of four selected candidate miRNAs (miR-16-5p, miR-29b-3p, miR-99a-5p, and miR-191-5p) which we identified by next-generation sequencing as being dysregulated. The selection of these miRNAs for further analysis by RT-qPCR was based on their degree of dysregulation (greater than five-fold change), availability of TaqMan™ Advanced miRNA assays, and expression abundance (>2,000 sequencing reads) to ensure detection sensitivity, experimental reproducibility, and biological significance. As with the sequencing data, the RT-qPCR data were analysed separately for males and females. To increase the statistical power of this analysis, we included an additional three samples per genotype per sex together with the original sequenced samples. The results are summarised in Table S5.

For miR-16-5p, the sequencing data indicated a significant effect of LPS treatment, compared to vehicle treatment, in female, but not male, samples, after adjusting for the genotype effects [false discovery rate (FDR)=0.0341, log_2_fold-change = −2.51]. Additionally, the sequencing data revealed a significant difference in the treatment response (i.e., difference in miR-16-5p expression between LPS and vehicle treatments) between female *TARDBP^M337V/M337V^* and female non-transgenic samples (FDR=0.0310, log_2_fold-change = 4.04). The normalised counts are shown in Fig. S1A-B. We did not see validation of the treatment effect of miR-16-5p in the RT-qPCR data (Fig. 6A). However, we validated the significant interaction between treatment and genotype (P=0.0206), for this miRNA. Post-hoc analysis of the simple genotype effects within each treatment level revealed this difference to be significant specifically between vehicle-treated *TARDBP^M337V/M337V^* and vehicle-treated *TARDBP^−/−^* samples (FDR=0.0119), but not between LPS-treated samples of the same genotypes (Fig. 6B).

**Figure 6.**
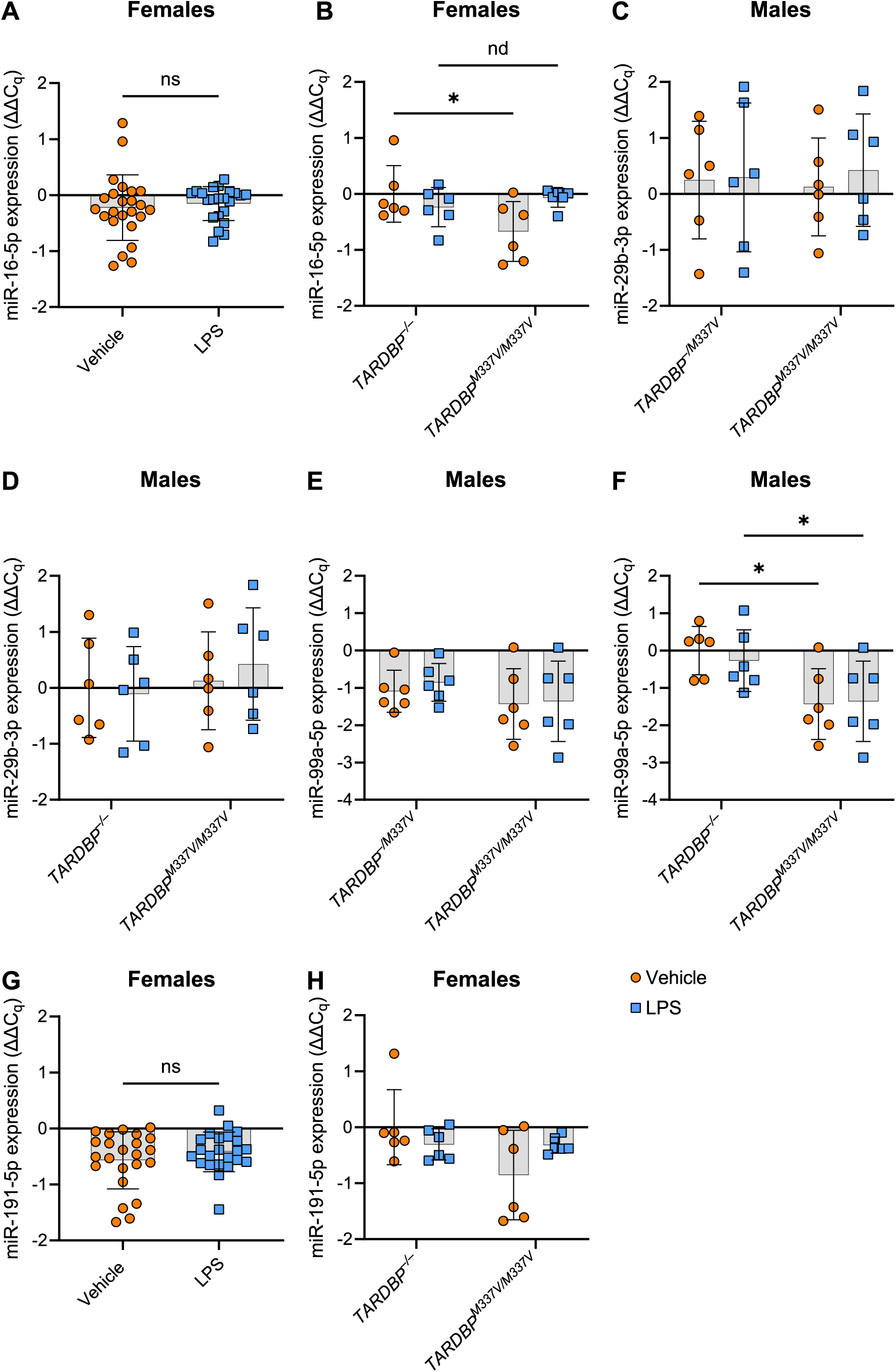
RT-qPCR validation results. Expression values (ΔΔCq) are normalised to the geometric mean expression of the vehicle-treated samples from *TARDBP^−/−^* animals within each sex group, following normalisation to three stable miRNAs (miR-28a-3p, miR-32-5p, and miR-190a-5p). Data is shown as mean ± SD. (**A**) Expression of miR-16-5p in female samples (all genotypes pooled). Paired two-tailed t-test. (**B**) Expression of miR-16-5p in female *TARDBP^−/−^* and *TARDBP^M337V/M337V^* samples. Repeated measures two-way ANOVA with two-stage step-up post-hoc. Significant genotype × treatment interaction (P=0.0206). (**C**) Expression of miR-29b-3p in male *TARDBP^−/M337V^* and *TARDBP^M337V/M337V^*samples. Repeated measures two-way ANOVA. No significant main effects. (**D**) Expression of miR-29b-3p in male *TARDBP^−/−^* and *TARDBP^M337V/M337V^*samples. Repeated measures two-way ANOVA. No significant main effects. (**E**) Expression of miR-99a-5p in male *TARDBP^−/M337V^* and *TARDBP^M337V/M337V^*samples. Repeated measures two-way ANOVA. No significant main effects. (**F**) Expression of miR-29b-3p in male *TARDBP^−/−^* and *TARDBP^M337V/M337V^*samples. Repeated measures two-way ANOVA with two-stage step-up post-hoc. Significant main effect of genotype (P=0.0250). (**G**) Expression of miR-191-5p in female samples (all genotypes pooled). Paired two-tailed t-test. (**H**) Expression of miR-29b-3p in female *TARDBP^−/−^* and *TARDBP^M337V/M337V^*samples. Repeated measures two-way ANOVA. No significant main effects. Panels **A** and **G**: N = 24 biological replicates. All other panels: N = 6 biological replicates per genotype. ns = not significant (P≥0.05), *discovery (FDR<0.05), nd = no discovery (FDR≥0.05).

For miR-29b-3p, the sequencing data showed a significant interaction between genotype and treatment when comparing male *TARDBP^M337V/M337V^*to male *TARDBP^−/M337V^* (FDR=0.0987, log_2_fold-change = 3.33), and when comparing male *TARDBP^M337V/M337V^* to male *TARDBP^−/−^* (FDR=0.0391, log_2_fold-change = 4.08). The normalised counts are shown in Fig. S1C-D. We did not see validation of these findings for miR-29b-3p in the RT-qPCR data (Fig. 6C-D). However, we found a statistically significant variation in miR-29b-3p expression between samples originating from different biological replicates (P=0.0071). This might have overshadowed any genotype or treatment effects in the RT-qPCR data.

For miR-99a-5p, the sequencing results identified a statistically significant difference in the treatment response between male *TARDBP^M337V/M337V^*and male *TARDBP^−/M337V^* samples (FDR=0.0460, log_2_fold-change = 4.18), as well as between male *TARDBP^M337V/M337V^* and male non-transgenic samples (FDR=0.0391, log_2_fold-change = 4.28). The normalised counts are shown in Fig. S1E-F. Analysis of the RT-qPCR data did not validate the differences between *TARDBP^M337V/M337V^* and *TARDBP^−/M337V^* samples (Fig. 6E). However, we found a statistically significant main effect of genotype (P=0.0250). Post-hoc analysis of the simple genotype effects within each treatment level revealed this difference to be significant specifically between vehicle-treated male *TARDBP^M337V/M337V^*and vehicle-treated male *TARDBP^−/−^* samples (FDR=0.0234), but also between LPS-treated *TARDBP^M337V/M337V^* and LPS-treated *TARDBP^−/−^* samples (FDR=0.0483), thus partially validating our sequencing findings (Fig. 6F). However, there was also a significant variation in miR-99a-5p expression between biological replicates (P=0.0025). This means that, even within the same genotype and treatment groups, samples from individual mice showed unique miR-99a-5p expression patterns.

For miR-191-5p, the sequencing data indicated a significant effect of LPS treatment, compared to vehicle treatment, in female, but not male, samples, averaged across all genotypes (FDR=0.00889, log_2_fold-change = −2.73). Additionally, the sequencing data revealed a statistically significant difference in the treatment response between female *TARDBP^M337V/M337V^*and female non-transgenic samples (FDR=0.0966, log_2_fold-change = 2.72). The normalised counts are shown in Fig. S1G-H. Analysis of the RT-qPCR data did not validate any of these findings for miR-191-5p (Fig. 6G-H).

Overall, our RT-qPCR analysis corroborated a significant miR-16-5p downregulation in female *TARDBP^M337V/M337V^*, compared to female *TARDBP^−/−^* samples, which is suppressed by LPS treatment. Also, a persistent downregulation of miR-99a-5p in male *TARDBP^M337V/M337V^*, compared to male *TARDBP^−/−^* samples, which persisted following LPS treatment, was observed. These findings partially validate our sequencing data using additional independent samples.

We then used the TargetScan online resource to predict the genes that our two validated dysregulated miRNAs target, followed by gene set enrichment analysis to determine whether any biological processes were overrepresented in the list of targets. This analysis revealed 195 predicted targets (Table S6), which were significantly enriched in biological processes related to neuronal function and development (Table S7), suggesting that miRNAs released from microglia have the potential to affect neighbouring neurons, provided they are internalised by them.

## Discussion

Our main aim was to determine whether there is a differential release of miRNAs from transgenic microglia, in the presence or absence of LPS stimulation. LPS is a known inducer of the transcription factor nuclear factor kappa-light-chain-enhancer of activated B cells (NF-κB). NF-κB regulates genes responsible for the innate and adaptive immune response, and several studies have shown that this transcription factor is upregulated in glial cells of both sporadic and familial ALS patients (Frakes et al., 2014, Maruyama et al., 2010, Swarup et al., 2011). The use of LPS as a stimulus has further relevance in the context of ALS, since previous research has shown elevated LPS levels in the blood plasma of sporadic ALS patients, which positively correlates with the levels of macrophage/monocyte activation (Zhang et al., 2009). In our study, stimulation of mouse microglia with LPS for 24 hours successfully resulted in a comparable level of pro-inflammatory response between males and females and between wildtype transgenic, mutant transgenic, and non-transgenic microglia, in terms of the expression of two inflammatory cytokines, *Tnf* and *Il1b*, at this timepoint. This suggests that the *TARDBP^M337V^* mutation does not alter the cytokine inflammatory response of microglia immediately after 24 hours of continuous exposure to LPS (at least for the tested cytokines). However, in the absence of LPS stimulation, the influence of the *TARDBP^M337V^* transgene on cytokine response appears to be complex since the baseline expression of *Tnf* was reduced by the presence of two copies of the mutant transgene in male microglia. Therefore, the precise effects of this mutation on the cytokine response of microglia, particularly in the context of different sexes, warrants further investigation.

A number of studies have observed that the intracellular miRNA content of microglia or monocytes is altered upon activation with different stimuli or in different disease states (Falcão et al., 2017, Butovsky et al., 2012, Krasemann et al., 2017). Furthermore, inflammation-related miRNA levels increase in the serum of mice following injection of LPS, although the source of these miRNAs was not determined (Li et al., 2018). The present study complements these findings by showing that, compared to vehicle treatment, LPS stimulation resulted in a significant alteration of certain miRNAs released by microglia, particularly in females. Previous research has already shown that male and female microglia exhibit gene expression differences and morphological differences, and that their response to environmental insults is sex-dependent (Dubbelaar et al., 2018, Thion et al., 2018, Hanamsagar et al., 2017). However, the present study is the first to observe sex-specific differences in terms of their released miRNAs. We also showed that the expression of the human *TARDBP^M337V^* transgene significantly affected the release of miRNAs from microglia, with some miRNAs showing a dose-dependent effect. These findings raise new questions about the role of mutant microglia in ALS. It may be that despite the apparently non-inflammatory phenotype of microglia in this TDP-43 mouse model, these cells can still influence the disease via miRNA pathways, especially given the known role of TDP-43 in RNA metabolism.

Furthermore, we confirmed the downregulated release of miR-16-5p and miR-99a-5p by homozygous mutant transgenic microglia. These miRNAs have been implicated in various biological processes, including cell proliferation, differentiation, apoptosis, cell cycle regulation, and the immune response (Cimmino et al., 2005, Liu et al., 2008, Bandi and Vassella, 2011, Rissland et al., 2011, Jing et al., 2005, Sun et al., 2011, Wang et al., 2017, Sun et al., 2014). There is a strong link between miR-16-5p and ALS in the literature, as we and others have shown that this miRNA is downregulated in the blood (Liguori et al., 2018, Joilin et al., 2020) and CSF (Waller et al., 2018) of sporadic ALS patients, as well as in fibroblasts from C9-ALS patients (Hur et al., 2023). Similarly, miR-99a-5p downregulation has also been observed in the muscle tissues from human ALS patients and the SOD1^G93A^ mouse model of ALS (Si et al., 2018), as well as in axons expressing TDP43^A315T^ and SOD1^G93A^ (Rotem et al., 2017). In light of these findings, the downregulated release of miR-16-5p and miR-99a-5p by homozygous mutant transgenic microglia may have significant implications in our understanding of ALS pathogenesis. Future studies should focus on elucidating the exact role these miRNAs play in ALS and whether modulating their levels could present a novel therapeutic approach.

We also conducted a parallel analysis for comparison purposes, utilising the same statistical procedures on the RT-qPCR data, but this time only including the original set of samples that were also subjected to RNA sequencing. This was carried out to assess the potential impact of sample size on the detection of the effects and interactions we were investigating. When we restricted our analysis to the original, smaller dataset, none of our results reached statistical significance. This does not necessarily discount the findings from the larger sample set, but rather highlights the sensitivity of such investigations to sample size. Inclusion of additional samples in our main analysis was intended as a strategy to increase the robustness of our findings and ensure they were not due to chance or random variability. These results underscore the importance of robust sample sizes in the detection of nuanced effects and interactions in biological systems, and the caution that should be exercised in interpreting findings from smaller sample sets.

Nevertheless, the discrepancies observed between the sequencing and RT-qPCR results could be attributed to several factors. First, technical variation between the two methods might contribute to the inconsistencies, due to differences in the efficiency of reverse transcription, amplification, and normalisation using reference miRNAs, which might have led to variations in the detected expression levels. Second, the addition of more samples in the RT-qPCR experiment may have introduced more biological variability. Finally, while RNA-seq has a broad dynamic range, RT-qPCR can often detect small changes in gene expression that are not detected by RNA sequencing.

Our gene set enrichment analysis revealed that the dysregulated miRNAs are predicted to target genes involved in neuronal development and function. It is important to experimentally validate some of these gene targets in future studies. Crucially, not only does a microglial miRNA need to find its way towards a neuronal cell, potentially via extracellular vesicles, but its target gene(s) must also be expressed within the neuron at that specific time, for it to have any functional effects. Nevertheless, we can hypothesise that the reduced release of our two candidate miRNAs and possibly others may affect normal neuronal function. Depending on whether the affected genes are positive or negative regulators of neuronal development and activity, this may implicate these miRNAs in neurodegeneration. Whilst we do not provide direct evidence that miRNAs released from microglia can have such effects on neurons, previous research has shown that uptake of miRNAs released from one cell type by another cell type (Aucher et al., 2013, Mittelbrunn et al., 2011, Zhou et al., 2014, Ridder et al., 2014, Pinto et al., 2017), including from microglia to neurons (Prada et al., 2018, Huang et al., 2017, Frühbeis et al., 2013, Varcianna et al., 2019), is possible. Furthermore, in addition to their gene repressor role, extracellular miRNAs have also been shown to activate intracellular neuronal receptors, leading to neurodegeneration (Park et al., 2014, Lehmann et al., 2012), or promoting recovery following neuronal insult (Xin et al., 2013), depending on which miRNAs are delivered. Since the microglia used in the current study were extracted from a mouse model of ALS, it would be interesting to investigate whether their released miRNAs can be taken up by disease-relevant cell types, such as cortical and spinal motor neurons, where they may target genes involved in neurodegeneration.

Traditionally, microglia have been implicated in neuroinflammatory diseases due to their reactive phenotype which is primarily characterised by cytokine release and phagocytosis, and several attempts have been made to develop treatments that modulate these processes. However, this study indicates that microglia should also be considered as possible regulators of gene silencing in other cell types via their miRNA release, thus opening new avenues in neuroinflammation research. In addition, TDP-43 mutations may further modulate this microglial response and thus have further implications in ALS research. Future studies could also examine whether other ALS-linked mutations, particularly mutations in genes involved in RNA metabolism, such as *FUS*, can similarly affect microglia-derived miRNAs.

## Materials and Methods

### Animals

Procedures involving animals were approved by the UK Home Office under the UK Animals Scientific Procedures Act 1986 and the University of Sussex Animal Welfare and Ethics Review Board. Mice (*Mus musculus*) were kept on a 12 h light/dark cycle (lights on at 7 am) in cages with up to five littermates, and free access to food and water. Transgenic mice carrying the human *TARDBP* gene (either wildtype or with the *M337V* mutation; JAX#029266) were bred with C57BL/6J mice (JAX#000664) to create in-house colonies. Generation of these transgenic mice is described elsewhere (Gordon et al., 2019). Briefly, a bacterial artificial chromosome (BAC) carrying the human *TARDBP* locus was integrated into the *Rosa26* locus of mouse embryonic stem cells which were then used to produce transgenic mice on a C57BL/6 background (Gordon et al., 2019). N = 12 mice (6 male and 6 female) per genotype (*TARDBP^−/−^*, *TARDBP^−/+^*, *TARDBP^−/M337V^*, *TARDBP^M337V/M337V^*) were used for primary mixed glial cell cultures. Due to difficulties encountered with the breeding and maintenance of *TARDBP^+/+^* mice (i.e., unexpectedly high number of unsuccessful breeders and overrepresentation of *TARDBP*^+/+^ mice that die on the first day after birth for reasons not investigated), the required number of homozygous *TARDBP*^+/+^ mice were not obtained, therefore they were not included in the study; however the hemizygous *TARDBP^−/+^* bred normally and were thus included in the study. Littermates were used wherever possible. The sex of all neonatal mice was determined by simplex PCR as previously described (Tunster, 2017).

### Coating of tissue culture surfaces

Cell culture flasks (75 cm^2^; Fisher, 15350591) were coated with 7 ml of poly-D-lysine (PDL; 10 μg ml^-1^ in molecular biology grade water), for 2 hours at 37°C. The flasks were then washed three times with 7 ml of molecular biology grade water and left to air-dry completely before use.

### Dissection and culture of primary mixed glia

Mixed glia were cultured as previously described (Lian et al., 2016), with some modifications. Briefly, the mice were sacrificed by cervical dislocation followed by exsanguination at postnatal day 0-3, and the brains were removed and kept immersed in dissection medium made up of Hank’s Balanced Salt Solution (HBSS++; Thermo Fisher Scientific, Gibco™, 24020117) containing 0.6% (w/v) D-glucose (Sigma-Aldrich, G-6152), 1% (v/v) of 1 M HEPES (Sigma-Aldrich, H-4034) at Ph 7.2–7.5, and 1% (v/v) Pen/Strep (Thermo Fisher Scientific, Gibco™, 15140122). Following removal of the meninges, the hippocampal and cortical tissue was minced using spring scissors and was transferred in a tube containing 30 ml cold dissection medium and 1.5 ml of 2.5% trypsin (Thermo Fisher Scientific, Gibco™, 27250018), in a 37°C water bath for 15 minutes, while swirling frequently, to dissociate the cells. Then, 1.2 ml of 1 mg ml^-1^ trypsin inhibitor [Sigma-Aldrich, T6522-25MG; suspended in Dulbecco’s Phosphate Buffered Saline (DPBS; Sigma-Aldrich, D8537-500ML)] was added to inhibit trypsin activity, and the tube was left to incubate for 1 minute at room temperature. After that, 750 μl of 1.2 mg ml^-1^ DNase I (Sigma-Aldrich, DN25-100MG; suspended in HBSS++) was added to digest the sticky DNA released from dead cells, and the tube was centrifuged at 400 × *g* for 5 minutes to pellet the cells. The supernatant was removed, the pellet was triturated in 5 ml pre-warmed culture medium containing 89% DMEM/F-12 (Fisher, Gibco™, 11580546), 10% heat-inactivated FBS (Fisher Scientific, Gibco™, 10270106), and 1% Pen/Strep (Fisher Scientific, Gibco™, 15140122), and centrifuged again at 400 × *g* for 5 minutes. The supernatant was removed, and the pellet was resuspended in 5 ml pre-warmed culture medium. The cell suspension was cultured in a PDL-coated flask with a total of 15 ml pre-warmed culture medium and incubated at 37°C, 5% CO2. The medium was replaced after 24 hours to remove cell debris. This culture consisted primarily of astrocytes and microglia (data not shown).

### Splitting flasks of mixed glia

After seven days in culture, each flask of mixed glia was split into two new flasks, to maximise the number of cells obtained per animal whilst minimising the number of animals used in accordance with the 3R principles of animal research. The culture medium was removed from the flask, and the cells were washed with 7 ml pre-warmed DPBS. Then, 2 ml of pre-warmed 0.25% trypsin-EDTA (Fisher Scientific, Gibco™, 11560626) was added to the cells and the flask was incubated for 5 minutes at 37°C, 5% CO_2_. After that, the flask was lightly tapped by hand to detach the cells, and 5 ml of pre-warmed culture medium was added to the flask to inactivate the trypsin. The medium with the cells was collected into a universal tube and centrifuged at 390 × *g* for 5 minutes to pellet the cells. The supernatant was removed, and the cells were resuspended with a P1000 micropipette in 5 ml of pre-warmed culture medium. The cell suspension was equally distributed into two new flasks, and the culture medium was topped up to 15 ml per flask. The flasks were then returned to 37°C, 5% CO_2_ for 14 days. Every 4-5 days, the culture medium of each flask was replaced. After approximately 7 days in culture, the flasks became 100% confluent, with a monolayer of astrocytes forming and microglia growing at the top and bottom of this monolayer.

### Isolation of microglia from mixed glial cultures

To remove astrocytes from the mixed cultures, a mild trypsinisation method resulting in >98% pure microglia cultures was used, as previously described (Saura et al., 2003), with some modifications. Briefly, the culture medium was removed, and the cells were washed with 7 ml pre-warmed DPBS. The mixed glia-conditioned medium was passed through a 0.2 μm filter to remove any cells and was kept for later. Then, 7 ml of 0.25% trypsin-EDTA diluted 1:4 in DMEM/F-12 was added to the cells and the flasks were incubated at 37°C, 5% CO_2_ for ∼1.5 hours, until the astrocyte monolayer and the microglia growing on top of it completely detached. The remaining, attached cells were only microglia. After that, 7 ml of pre-warmed culture medium was added to the flask to inactivate the trypsin, and the entire medium with the floating cells was discarded. The remaining microglial cells were washed twice with 7 ml pre-warmed DPBS to ensure the removal of astrocytes. Then, 4 ml of 0.25% trypsin-EDTA was added to the flask, and the cells were incubated for 5 minutes at 37°C, 5% CO_2_ to detach them. After incubation, the cells were scraped off the bottom of the flask using a cell scraper, and then 8 ml of pre-warmed culture medium was added to inactivate the trypsin. The cell suspension from both flasks of each animal was collected into a single tube and centrifuged for 10 minutes at 390 × *g* to pellet the cells. The supernatant was removed, and the cells were resuspended in 2 ml of astrocyte-conditioned medium (this was a mixture of 50% fresh culture medium and 50% mixed glia-conditioned culture medium collected at the beginning of the isolation). The cells were counted with a haemocytometer and plated at 100,000 cells/well in 24-well plates (1 ml/well), using 50% fresh and 50% conditioned medium, and incubated at 37°C, 5% CO_2_ for 48 hours. The purity of the microglia cultures was estimated by microscopic inspection to be >98% (data not shown).

### Lipopolysaccharide (LPS) stimulation of microglia in culture

Forty-eight hours following plating of the microglia, the culture medium was discarded, the cells were washed with 1 ml/well of pre-warmed DPBS to ensure complete removal of remaining serum, and 1 ml/well of pre-warmed serum-free culture medium was added to the cells for a further 24 hours. This medium change was necessary to remove any molecules released by the cells which are triggered by the isolation procedure or induced by the attachment to the plate, as these could potentially influence the inflammatory state of the microglia. Furthermore, serum may contain proteins that act as cofactors for various agonists and activate undesired receptor systems that could lead to responses unrelated to what is being measured, so its removal is necessary to avoid this. Importantly, serum has been shown to contain small ncRNA contaminants (Mannerström et al., 2019), which could confound the detection of microglia-derived ncRNAs in subsequent experiments.

Following a rest period of 24 hours in serum-free culture medium, the cells were treated with fresh serum-free culture medium containing either 250 ng ml^-1^ LPS (Sigma-Aldrich, L2654-1MG) or vehicle (HBSS), for a further 24 hours. The LPS concentration and incubation time were selected based on optimisation experiments showing 24 hours is sufficient to substantially stimulate the cells without significant cytotoxicity at this concentration (data not shown). Furthermore, optimisation experiments also suggested that 24 hours is sufficient time for extracellular miRNAs to accumulate into the culture medium in the absence of any stimulation (data not shown). After stimulation, the culture medium was collected and immediately frozen at −80°C. The cells were washed with 1 ml/well pre-warmed DPBS, then 260 μl of buffer RLT (Qiagen, 79216) containing 1% β-mercaptoethanol (Fisher Scientific, M/P200/05) was added to the cells to detach and lyse them. The cell suspension was collected into a tube and immediately vortexed for 1 minute to homogenise the cells, then snap-frozen in liquid nitrogen, followed by long-term storage at −80°C until RNA extraction.

### Intracellular RNA extraction, cDNA synthesis, and RT-qPCR

The total RNA from the collected microglia was extracted using the miRNeasy Tissue/Cells Advanced Mini Kit (Qiagen, 217604), according to the manufacturer’s protocol. The amount of RNA was quantified using the Qubit™ RNA high sensitivity assay kit with the Qubit™ 3.0 fluorometer. Complementary DNA (cDNA) synthesis using 8 ng of total RNA was carried out using the High-Capacity RNA-to-cDNA kit (Thermo Fisher Scientific, 4388950), according to the manufacturer’s protocol. A 10-fold dilution of the cDNA was then used for SYBR-Green RT-qPCR to quantify the expression levels of *Tnf* and *Il1b* (normalised to the expression levels of *Gapdh* and *Pgk1*), using the SYBR qPCR 2x mix (AptoGen, 411101.5.1A25) and the following primers (5’-to-3’ sequence): *Tnf* forward primer, GGT GCC TAT GTC TCA GCC TCT T; *Tnf* reverse primer, GCC ATA GAA CTG ATG AGA GGG AG; *Il1b* forward primer, TGC CAC CTT TTG ACA GTG ATG; *Il1b* reverse primer, TGA TGT GCT GCT GCG AGA TT; *Gapdh* forward primer, GGT GAA GGT CGG TGT GAA CG; *Gapdh* reverse primer, CAA TCT CCA CTT TGC CAC TGC; *Pgk1* forward primer, TTG TGC ATT GTA GAG GGC GT; *Pgk1* reverse primer, TGA CGA AGC TAA CCA GAG GC.

### Extracellular miRNA extraction, miRNA library preparation, Illumina sequencing, cDNA synthesis, and TaqMan™ advanced miRNA assays

Extracellular RNA was extracted from 200 μl of microglia-conditioned culture medium using the miRNeasy Serum/Plasma Advanced Kit (Qiagen, 217204). The QIAseq miRNA Library Kit (Qiagen, 331505) was used with 5 μl of extracellular RNA to prepare 48 miRNA libraries for next generation sequencing according to the manufacturer’s protocol, but with the following modifications: (1) the 3’ adapter and the RT primer were used at a 1:20 dilution, (2) the 5’ adapter was used at a 1:10 dilution, (3) all bead washing steps were done using 500 μl of 80% instead of the recommended 200 μl, and (4) at the library amplification step, denaturation, annealing, and extension were performed for 24 cycles. The tube indices from the QIAseq miRNA NGS 48 Index IL (Qiagen, 331595) were used for library preparation.

Pre-sequencing quality control of the miRNA libraries was done on an Agilent Bioanalyzer using a High Sensitivity DNA chip (Agilent Technologies, 5067-4626). Then, the Qubit™ dsDNA Assay Kit (Thermo Fisher Scientific, Q32854) was used with the Qubit™ 3.0 Fluorometer to quantify the amount of library before diluting and pooling the samples at equimolar concentrations for sequencing. The sequencing of the miRNA libraries was performed by the next-generation sequencing facility at the University of Leeds, where the BluePippin was used to size-select for the libraries between 170 and 200 bp, to extract miRNA and piRNA libraries. The Bioanalyzer and Qubit™ were then used to confirm the presence of correctly sized libraries and quantify the amounts. The libraries were then prepared for 100 bp single-end sequencing using the P3 kit on the Illumina NextSeq 2000, with each sample split over two lanes on a flow cell. Upon completion of the sequencing, the sequencing facility performed the basecalling and trimmed the 5’ adapters from the reads.

Furthermore, 2 μl of extracellular RNA were used with the TaqMan™ Advanced miRNA cDNA Synthesis Kit (Thermo Fisher Scientific, A28007). The miRNAs with the least expression variability between the 48 sequenced samples were identified using geNorm (Vandesompele et al., 2002) in R version 4.2.0, and the three least variable miRNAs were selected as normalisers for RT-qPCR validation. Specific miRNAs of interest were then used with the following TaqMan™ Advanced miRNA assays (Fisher, 15412184) to validate the sequencing results by RT-qPCR, according to the manufacturer’s protocol, using fast-cycling and with the qPCR Lo-ROX 2X (AptoGen, 412101.5) mix. The TaqMan™ Advanced miRNA assays for the normaliser miRNAs were mmu-miR-28a-3p (mmu481665_mir), mmu-miR-32-5p (mmu482950_mir), and mmu-miR-190a-5p (mmu481335_mir). Those for the target miRNAs were mmu-miR-16-5p (mmu482960_mir), mmu-miR-29b-3p (mmu481300_mir), mmu-miR-99a-5p (mmu478519_mir), and mmu-miR-191-5p (mmu481584_mir).

### Bioinformatic analysis

Quality control of the sequencing data was undertaken using the FastQC tool (Andrews, n.d.) via the Galaxy webserver (Afgan et al., 2018). Primary miRNA quantification was done using predefined analysis pipelines via the QIAGEN RNA-seq Analysis Portal 2.0 (available at ngsdataanalysis2.qiagen.com/QIAseqmiRNA). The reads were aligned to the latest mouse genome assembly available at the time of analysis (GRCm38.p6; RefSeq assembly accession GCF_000001635.26) using Bowtie (Langmead et al., 2009). This alignment strategy mapped the reads to those available in miRBase V21 (Kozomara et al., 2019). Secondary analysis to calculate differential miRNA expression between the different samples was undertaken in R (version 4.2.0) (R Core Team, 2020) using the DESeq2 package (release 3.15) (Love et al., 2014). We designed our analysis to consider both the effects of genotype and treatment on miRNA expression, as well as the potential interaction between these two factors.

Specifically, our design formula was ∼ Mouse + Genotype + Treatment + Genotype:Treatment. The “Mouse” term in the model accounts for the pairing of samples from each biological replicate, adjusting for their inherent non-independence due to one sample receiving LPS treatment and the other receiving the vehicle treatment. Sex was excluded from the model to maintain simplicity and avoid overfitting. Therefore, the model was applied to each sex separately to address potential sex-specific effects. We then performed a series of contrasts to understand the different effects in our experiment. Firstly, the effect of treatment was assessed by comparing miRNA expression between the LPS and Vehicle conditions. In other words, this comparison answers the question “How does the miRNA expression for the LPS treatment compare to the Vehicle treatment, after controlling for the genotype effect?”. Secondly, to examine the effect of different genotypes, we performed three comparisons: (1) *TARDBP^M337V/M337V^* versus *TARDBP^−/−^*, (2) *TARDBP^−/M337V^*versus *TARDBP^−/−^*, and (3) *TARDBP^−/+^* versus *TARDBP^−/−^*. In other words, each of these comparisons answers the question “How does the miRNA expression in Genotype X compare to the control (*TARDBP^−/−^*) genotype, after controlling for the treatment effect?”. Finally, we performed pairwise contrasts of the interaction terms to understand how the response to LPS treatment varies between different genotypes. Specifically, we examined the differences between (1) *TARDBP^M337V/M337V^* and *TARDBP^−/−^*, (2) *TARDBP^M337V/M337V^*and *TARDBP^−/M337V^*, (3) *TARDBP^M337V/M337V^* and *TARDBP^−/+^*, (4) *TARDBP^−/M337V^* and *TARDBP^−/+^*, (5) *TARDBP^−/M337V^* and *TARDBP^−/−^*, and (6) *TARDBP^−/+^*and *TARDBP^−/−^*. In other words, each of these comparisons answers the question “How does the difference in miRNA expression between LPS and Vehicle treatments for Genotype X compare to that same difference for Genotype Y?”. Any miRNAs with a false discovery rate (FDR) <10% were considered to be significantly dysregulated. Target prediction of candidate miRNAs was performed using the TargetScan online resource (version 8.0) (Agarwal et al., 2015), and gene set enrichment analysis of the predicted target genes of the candidate miRNAs was carried out using the gene ontology online resource (Ashburner et al., 2000, Aleksander et al., 2023).

### Statistical analysis

RT-qPCR data were analysed in GraphPad Prism version 10.0.0. Outliers were identified by the ROUT test with Q = 1% and were removed before further analysis, as indicated in the figure legends. Data on *Tnf* and *Il1b* mRNA expression were analysed using a two-way repeated measures analysis of variance (ANOVA) with Šidák’s post-hoc test. Data on miRNA expression were analysed with either paired two-tailed t-tests or with two-way repeated measures ANOVA tests, as indicated in the figure legends. The threshold for significance was set to 5%. Where statistically significant main effects were identified by the ANOVA tests, post-hoc analysis was performed using the two-stage step-up method of Benjamini, Krieger and Yekutieli to correct for multiple comparisons by controlling the FDR (set to 5%).

## Acknowledgements

We would like to thank the Next Generation Sequencing Facility at the University of Leeds for the generation and initial processing of the sequencing data.

## Data availability

The RNA sequencing data associated with this article are publicly available through the European Nucleotide Archive under the project accession number PRJEB61796.

## Supplementary materials

**Figure S1.**
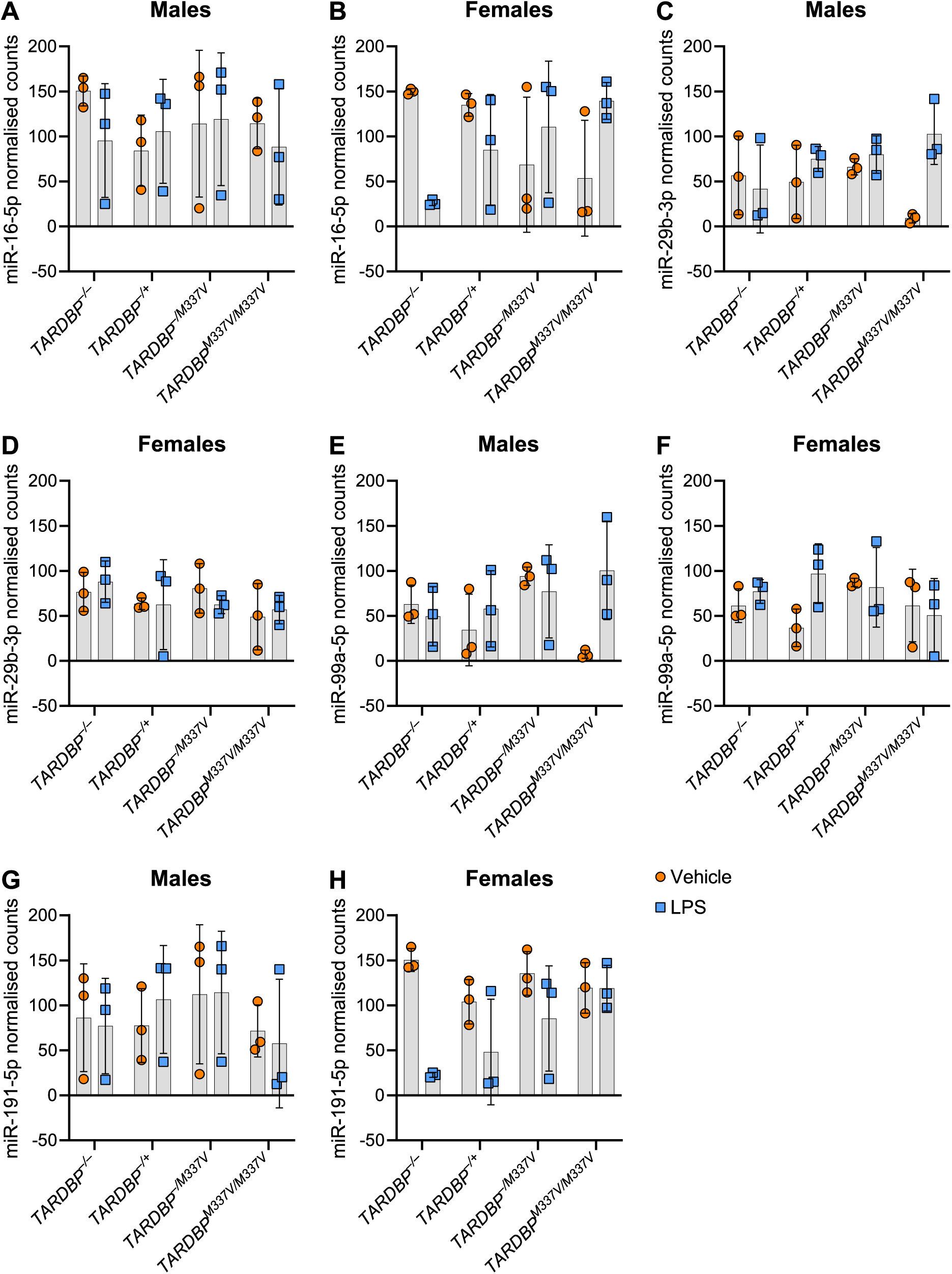
Normalised counts used by DESeq2 for analysis, for the four candidate miRNAs selected for RT-qPCR validation. (**A-B**) Normalised counts for miR-16-5p. (**C-D**) Normalised counts for miR-29b-3p. (**E-F**) Normalised counts for miR-99a-5p. (**G-H**) Normalised counts for miR-191-5p. Data is shown as mean ± SD. *N* = 3 biological replicates per genotype per sex.

**Table S1**. Dysregulated release of miRNAs upon LPS stimulation, after adjusting for genotype effects.

**Table S2**. Dysregulated release of miRNAs in female transgenic samples, after adjusting for treatment effects.

**Table S3**. Interaction between genotype and treatment in dysregulating release of miRNAs.

**Table S4**. Overlap between different genotypes in terms of the differences in the response to LPS treatment, compared to vehicle treatment.

**Table S5.** Summary of statistically significant results in the DESeq2 data and corresponding results in the RT-qPCR data.

**Table S6**. Predicted targets of mmu-miR-16-5p and mmu-miR-99a-5p.

**Table S7**. Biological processes significantly enriched in the list of predicted targets of mmu-miR-16-5p and mmu-miR-99a-5p.

